# Global Prevalence of *Cryptosporidium* Infections in Cattle and *C. parvum genotype* distribution: A Meta-Analysis

**DOI:** 10.1101/2024.07.16.603704

**Authors:** Rachel Buchanan, Eleni Matechou, Frank Katzer, Anastasios D. Tsaousis, Marta Farré

**Affiliations:** School of Biosciences, University of Kent; School of Mathematics, Statistics and Actuarial Sciences, University of Kent; Moredun Research Institute, Scotland, UK

**Keywords:** *Cryptosporidium*, Cryptosporidiosis, Cattle, metanalysis, epidemiology

## Abstract

**Background:** The protozoan parasite *Cryptosporidium* is the causative agent of a severe diarrhoeal disease, called cryptosporidiosis. *Cryptosporidium* species are capable of infecting a wide range of hosts including humans and livestock. In cattle, cryptosporidiosis is now one of the most important causes of neonatal scour globally, either as a sole agent or co-infecting with other pathogens. Cryptosporidiosis is considered globally endemic, with a prevalence of *Cryptosporidium* in stool samples from 13% to 93% in European cattle. This disease has a significant economic burden, with costs associated with veterinary diagnosis, medication, increased labour, animal rearing and supplemental nutrition as well as being associated with reduced long-term growth rate in calves, causing huge economic losses in livestock industry. Moreover, cattle act as a zoonotic reservoir for *Cryptosporidium parvum*, a species that is capable of infecting humans as well. As such, monitoring the prevalence of *Cryptosporidium* in cattle is important due to the public health risk and financial burden the clinical disease causes.

**Methods:** Publications reporting on the prevalence of *Cryptosporidium* in cattle were collected from PubMed and Google Scholar. Information regarding the species of *Cryptosporidium* in positive samples, the genotype of *C. parvum* found in samples, and the diarrhoeic status of the cattle was collected where available. A total of 279 publications were collected for this meta-analysis from six continents and 65 countries to provide an estimation for global bovine *Cryptosporidium* prevalence.

**Results:** A 25.5% global prevalence of *Cryptosporidium* infection was reported, with *C. parvum* being the most frequently identified species, particularly the IIa subfamily. Diarrhoea was reported in 14,141 cattle samples, of which 36.0% tested positive for *Cryptosporidium*. Regarding symptoms, we found that in countries reporting over 50% of diarrhoeic positive cattle, *C. parvum* was the most common species.

**Conclusions:** Continued monitoring and reporting of *Cryptosporidium* in cattle are crucial for both public health and economic reasons. Consequently, efforts should focus on underreported regions and the development of control measures to reduce prevalence and limit zoonotic transmission.

## BACKGROUND

Since its identification in mice in 1907 [1] there are now more than 40 recognised species of the apicomplexan parasite *Cryptosporidium* [2], the causative agent of the diarrhoeal disease cryptosporidiosis. Of these species, four are common in cattle: *C. parvum*, *C. bovis*, *C. andersoni*, and *C. ryanae* [3–5]. There is an age-related distribution of the common species, with *C. parvum* being the predominant species in pre-weaned dairy calves, *C. bovis* and *C. ryanae* in post-weaned dairy calves, and *C. andersoni* in yearling calves and adults [6,7]. It is likely that the same distribution exists in beef cattle [8,9].

While four species are commonly found in cattle, only *C. parvum* and *C. andersoni* have been associated with clinical disease, and thus can financially burden cattle farms. *C. parvum* infection frequently leads to diarrhoeal disease in pre-weaned calves [10,11], and *C. andersoni* has been associated with a reduction in milk production and weight loss in adult cattle [12,13]. Genotyping of *Cryptosporidium* at the *60kDa glycoprotein* (*gp60*), *heat shock protein 70 (Hsp70), actin* and *COWP* has resulted in the identification of over 120 different genotypes [2]. In cattle, three subfamilies have been identified: IIa, IId, and IIl. The IIa subfamily, which includes the highly transmissible IIaA15G2R1 genotype [14], is most frequently found in Europe, while the IId subfamily is more prevalent in Asia, especially in China [15].

*Cryptosporidium* is a zoonotic pathogen which occurs in cattle, and was the second most frequent diagnosis in diseased cattle between 2015 and 2023 in the Great Britain [16]. Rises in the global number of Concentrated Animal Feeding Operations (CAFOs), particularly in industrialised countries have been associated with increased transmission of several pathogens including *C. parvum,* posing a further risk for increased transmission to humans [17].

As the global demand for livestock products, including cattle, is expected to increase in developing countries with a growing population [23], monitoring the global distribution of *Cryptosporidium* in cattle is important due to the public health risk of transmission and for the health of farmed animals. Two previous meta-analyses have given estimations of global *Cryptosporidium* infections in cattle [24,25]. In 2019, Hatam-Nahavandi et al. presented the prevalence of *Cryptosporidium spp.* in terrestrial ungulates including cattle, and in 2023 Chen et al. presented the prevalence of *C. parvum* genotypes in dairy calves. Here, we expand on previous publications by first conducting a systematic review and meta-analysis of the global prevalence of *Cryptosporidium spp.* including *C. parvum* genotypes in cattle of all age groups. We analysed the health status of the animals and discuss the link between *C. parvum* and the prevalence of diarrhoea during infection. By not restricting the characteristics of sampled cattle, we present data for a larger sample size of 144,523 cattle to provide an updated estimation of global species and *C. parvum* genotype distribution.

## Methods

### Search strategy

To identify publications reporting on *Cryptosporidium* prevalence in cattle, PubMed and manual searching via Google Scholar were used. Only papers published between 2003 and October 2023 were included in the meta-analysis. In Google Scholar, searches were made using combinations of key words ‘Cattle’, ‘Calf’, ‘Calves’, *‘Cryptosporidium’*, and ‘Cryptosporidiosis’. In PubMed, publications were identified using MeSH terms with the formula (((((cattle[MeSH Terms])) OR (calf)) OR (calves)) AND (cryptosporidium[MeSH Terms])) OR (cryptosporidiosis[MeSH Terms])).

### Publication inclusion and exclusion criteria

Publications were initially excluded based on title and abstract screening for relevance. Publications identified as potentially relevant were read in full and were included in the meta-analysis if the inclusion criteria were met. Publications were excluded based on several factors: incorrect host species, publication was a review article, the full text was not available or not in English, the prevalence of *Cryptosporidium* was not stated, or prevalence results in the publication were already published in a different publication.

### Data extraction

Data was extracted from publications which met the inclusion criteria. The following data was collected: publication year, first author, location of study (country), host species, clinical signs (diarrhoeic or non-diarrhoeic), age(s) of sampled cattle, detection method used, keeping status, total samples collected, total samples tested positive for *Cryptosporidium*, *Cryptosporidium* species identified, and *C. parvum gp60* genotypes identified.

### Meta-analysis

The meta-analysis was conducted with the aim to enhance knowledge of global *Cryptosporidium* prevalence and the distribution of species and *C. parvum* gp60 genotypes, given the burden of cattle losses to cryptosporidiosis and considering that cattle play a role in zoonotic transmission. All data was analysed in RStudio (Version 4.3.1). The R packages meta (Version 6.5-0) and metafor (Version 4.4-0) were used to calculate the pooled prevalence of *Cryptosporidium* infection in cattle along with the 95% confidence interval. Due to high heterogeneity (*I^2^* > 50%, p < 0.1) a random effects model was used for meta-analysis. Forest plots were generated to display results by continent and by method used for *Cryptosporidium* detection. If multiple methods were used for *Cryptosporidium* detection in the same samples, the molecular method was included in analysis. The extent of publication bias was also tested using a funnel plot and Egger’s test. Sources of heterogeneity were assessed through meta-regression using the R package metafor (Version 4.4-0). This meta-analysis and literature review are conducted in accordance with the PRISMA guidelines [26] (Table S1).

## Results

Database searching revealed 1,629 potentially relevant publications, of which 1,262 were removed during initial screening of the publication titles and abstracts. The remaining 367 publications were further assessed, with 88 publications excluded due to not meeting the inclusion criteria. After all publications were assessed, 279 publications met the inclusion criteria and were included in the meta-analysis. These 279 publications resulted in 292 studies for use in the meta-analysis (Figure 1a, Table S2). This includes a total of 41 studies from Africa, 119 studies from Asia, 71 studies from Europe, 26 studies from North America, 15 studies from Oceania, and 20 studies from South America (Figure 1b and Additional File 2: Table S2).

**Figure 1.**
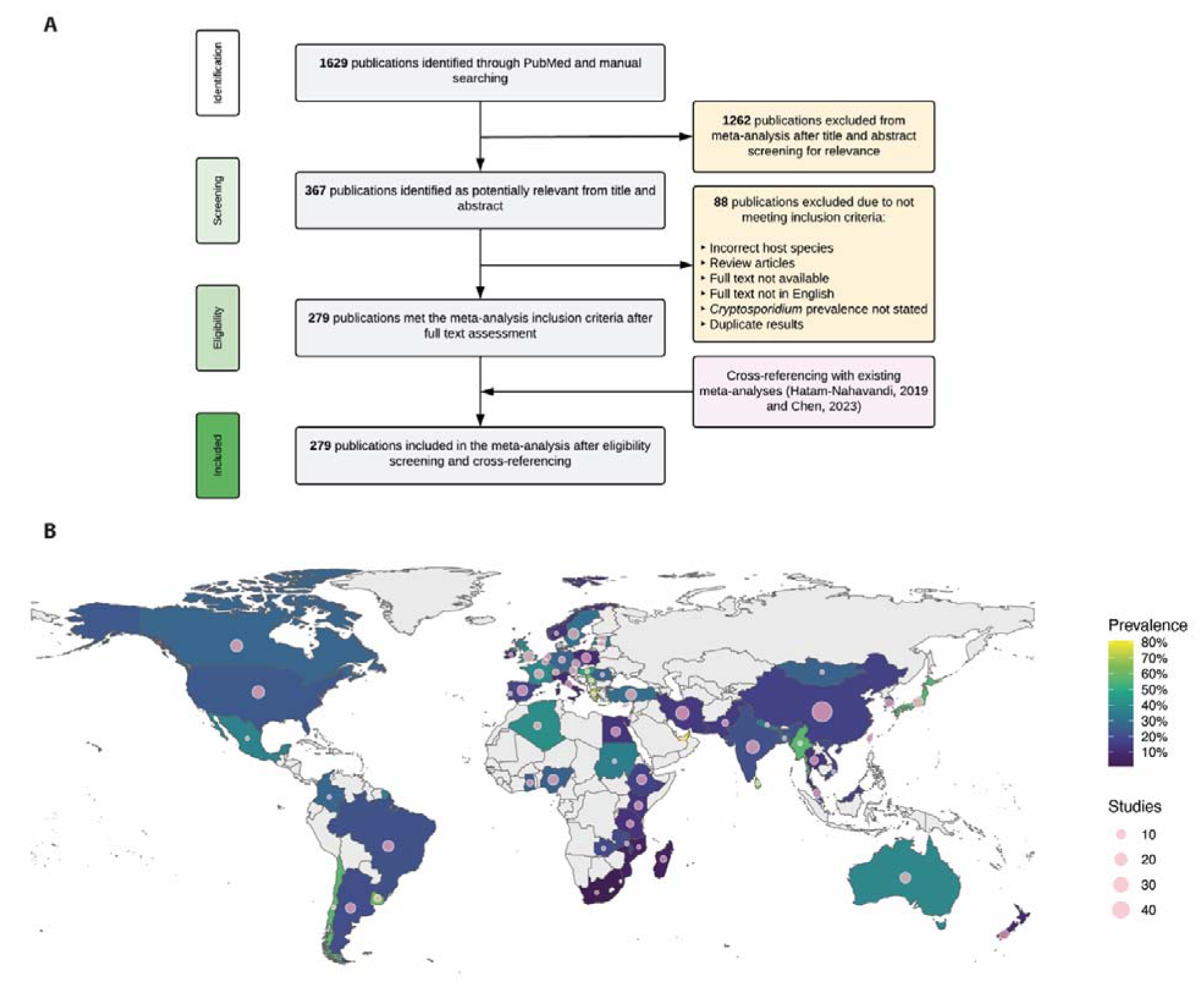
Global prevalence of *Cryptosporidium*. **A.** Flow diagram of the publication inclusion process. **B.** Estimation of bovine *Cryptosporidium* prevalence and the number of studies per country.

The overall pooled prevalence of *Cryptosporidium* worldwide was 25.5% [95% CI: 23.3-27.7%], comprising of samples from 144,523 cattle. This prevalence ranges from 20.0% [95% CI: 16-24%] in Africa to 30.5% [95% CI: 26.0-35.0%] in Europe (Figure 2), with South America and Oceania presenting similar prevalence (27.1% [95% CI: 15.8-38.4%] and 27.2% [95% CI: 18.1-36.4%], respectively (Figure 2).

**Figure 2.**
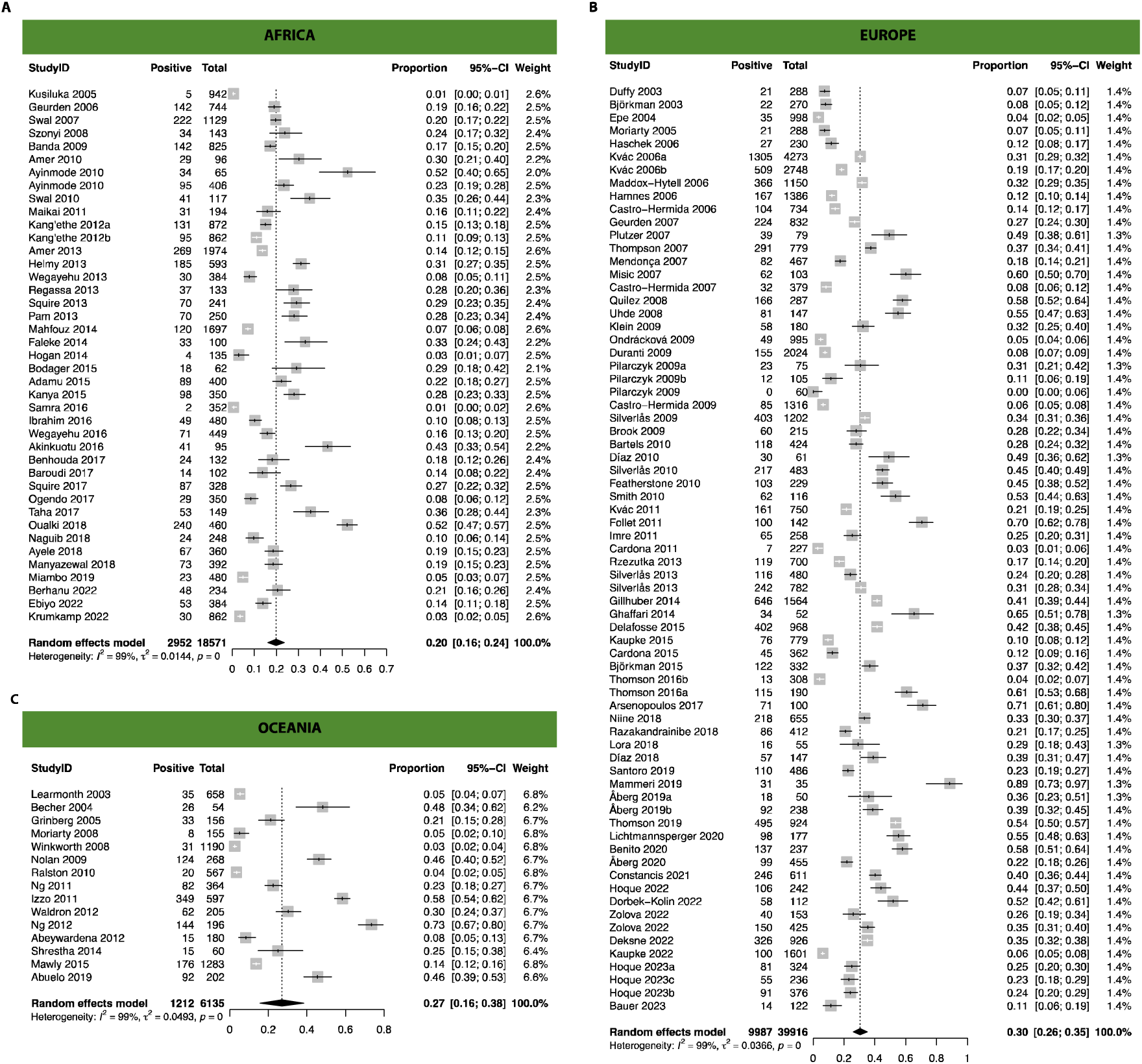
Forest plots of estimated *Cryptosporidium* prevalence in cattle from studies conducted in a) Africa, b) Europe, and c) Oceania.

We identified studies comprising 65 countries, with one study in 26 countries, 2-10 studies in 34 countries and only five countries included data from more than 10 papers (Figure 1b). The prevalence of the majority of countries falls between 10-30%. The country with the lowest prevalence was South Africa (0.6%), however this was only based on one study. Similarly, the country with the highest prevalence was the United Arab Emirates (80.6%), however this was again based on one study. Japan, with eight studies, has a prevalence of 54.7%, giving it the highest prevalence of countries with more than five studies. New Zealand, with seven studies, has a prevalence of 8.5%, the lowest of all countries with more than five studies. This low reported prevalence in New Zealand was also identified in a previous meta-analysis [24].

*Cryptosporidium* can be detected using several methods, including conventional microscopy (CM), immunochromatographic test (ICT), immunofluorescence antibody test (IFA), PCR and ELISA, all with different detection capacities. To assess whether the method of detection used in each study might impact the estimated pooled prevalence, we grouped the studies by method of detection and calculated the pooled prevalence in each type of method used. The ELISA method resulted in a higher pooled prevalence (43.2% [95% CI: 28.3-58.1%]) than the other detection methods, while the pooled prevalence of the most used methods (CM and PCR) was 25.0%, with 95% CI: 21.9-28.1% and 95% CI: 21.2-28.7%, respectively (Table 1). However, the high pooled prevalence for studies using ELISA might have resulted due to the study targeting only symptomatic cattle, as only 6 of the 16 studies utilising the ELISA method reported on the diarrhoeic status of the sampled cattle.

**Table 1.** Summary of bovine *Cryptosporidium* prevalence and statistical analysis by continent, detection method and keeping status.

|  | Pooled Prevalence |  | Heterogeneity |  |  | Publication bias |  |  |
| --- | --- | --- | --- | --- | --- | --- | --- | --- |
|  | Pooled (%) | OR (95% CI) | *I*<sup>2</sup> (%) | Q | df | Egger bias | t-value | Egger p-value |
| Global | 25.5 | 23.3-27.7 | 99.4 | 48174.7 | 291 | 11.9 | 12.4 | p < 0.0001 |
| **Continent** |  |  |  |  |  |  |  |  |
| Africa | 20.0 | 16.1-23.7 | 98.5 | 2723.6 | 40 | 9.8 | 10.5 | p < 0.0001 |
| Asia | 23.9 | 20.4-27.5 | 99.5 | 24074.0 | 118 | 11.3 | 6.4 | p < 0.0001 |
| Europe | 30.5 | 26.0-35.0 | 99.0 | 6908.2 | 70 | 11.3 | 7.0 | p < 0.0001 |
| North America | 25.2 | 16.4-33.9 | 99.7 | 7320.9 | 25 | 13.9 | 2.6 | p = 0.0152 |
| Oceania | 27.1 | 15.8-38.4 | 99.2 | 1698.3 | 14 | 12.4 | 4.1 | p = 0.0014 |
| South America | 27.2 | 18.1-36.4 | 99.1 | 2126.1 | 19 | 13.6 | 2.6 | p = 0.0189 |
| **Detection Method** |  |  |  |  |  |  |  |  |
| CM | 25.0 | 21.9-28.1 | 99.3 | 17910.6 | 122 | 12.3 | 9.8 | p < 0.0001 |
| ELISA | 43.2 | 28.3-58.1 | 99.9 | 10978.13 | 15 | 7.1 | 0.6 | p = 0.5412 |
| ICT | 21.3 | 13.2-29.4 | 88.6 | 43.9 | 5 | **Too small (6 studies)** |  |  |
| IFA | 23.1 | 17.5-28.9 | 99.1 | 4168.4 | 39 | 12.1 | 5.6 | p < 0.0001 |
| PCR | 25.0 | 21.2-28.7 | 99.0 | 10945.9 | 104 | 11.5 | 11.9 | p < 0.0001 |
| **Keeping Status** |  |  |  |  |  |  |  |  |
| Beef | 20.1 | 11.4-30.2 | 99.0 | 1142.2 | 12 | 12.3 | 4.0 | p = 0.0020 |
| Dairy | 28.0 | 24.5-31.3 | 99.3 | 20210.4 | 134 | 12.4 | 8.9 | p < 0.0001 |

*Cryptosporidium* usually leads to diarrhoea in most animals infected; however, in some instances infected animals do not develop symptoms. To estimate the proportion of these cases around the globe, we analysed a total of 88 studies across 36 countries reporting on the prevalence of diarrhoea in the sampled cattle (Figure 3a). Of these studies, 18 countries only had data from one study reporting on diarrhoea prevalence, 17 countries had 2-8 studies, while 13 studies in China reported this information. Two countries (Mozambique and Tanzania) reported positive cattle that were not diarrhoeic, while the majority of countries reported over 50% of positive cattle exhibiting diarrhoea (Figure 3a). In all these countries, *C. parvum* was the highest reported species, except for Bangladesh, where there was no species data. Interestingly, China and the Czech Republic are two examples with larger available datasets where less than 50% of cattle exhibited diarrhoea and *C. parvum* was not the dominant species. Because diarrhoea can be caused by other infections, 23 studies analysed the presence of *Cryptosporidium* in these animals (Fig 3b). An average of 36% of cattle with diarrhoea tested positive across all countries, but a large variation between countries was observed, with Egypt and Republic of Korea reporting the lowest percentages of 9.2% and 11.9%, respectively, while Uruguay and France had the highest percentages, with 69.6% and 72.0%, respectively.

**Figure 3.**
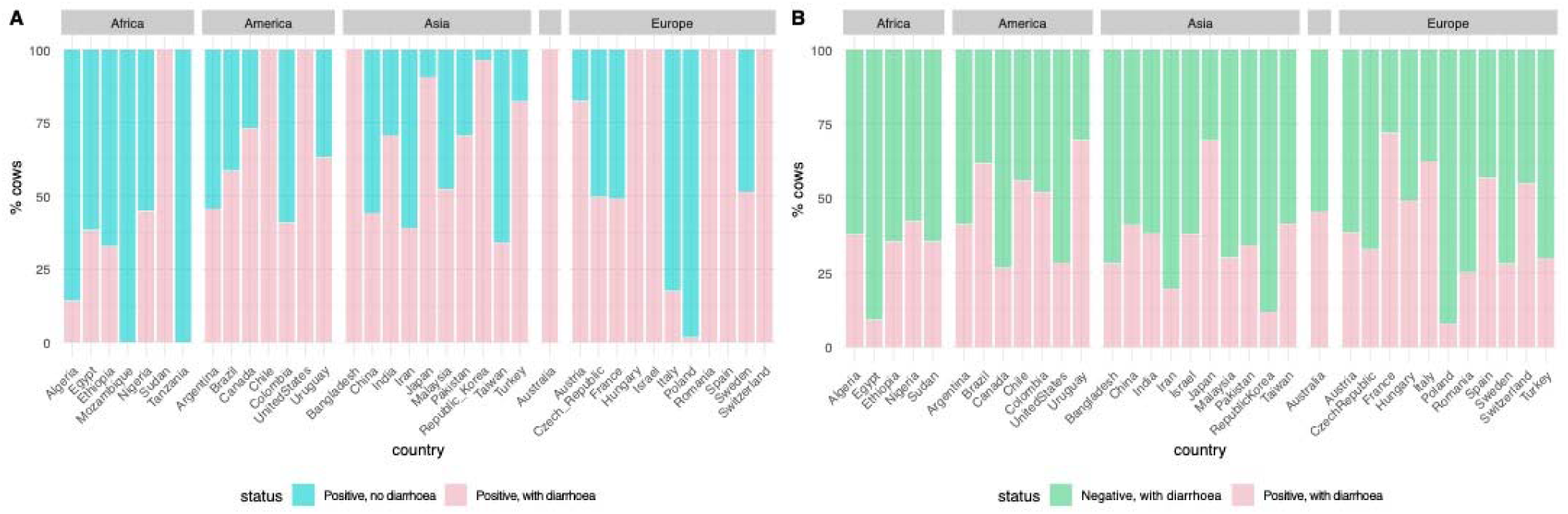
Global estimates of a) infected cattle with or without diarrhoea and b) diarrhoeic cattle that are either positive or negative for *Cryptosporidium* infection.

A total of 65.7% (20,520/31,239) of animals infected by *Cryptosporidium* were assessed to determine the exact species causing the infection. The number of samples with detailed *Cryptosporidium* species ranged from 88.6%, 74.1%, 72.3%, 62.4%, 34.5%, and 31.3% of samples in Oceania, Europe, Asia, North America, South America, and Africa, respectively. By country, this percentage varied, with several countries’ studies determining all or more than 75% of samples, while other countries’ studies determined none or less than 25% of samples. From all studies, 98.5% (20,217/20,520) were mono-infections, with 303 reported incidents of mixed *Cryptosporidium* spp infections, the majority of which were identified as *C. bovis* + *C. ryanae* (131 cases) or *C. bovis* + *C. parvum* (83 cases). Mono-infection of *C. parvum* was identified in 61.7% (12,664/20,520) of samples, *C. bovis* in 15.3% (3146/20,520) of samples, *C. andersoni* in 14.4% (2961/20,520) of samples, and *C. ryanae* in 5.6% (1152/20,520) of samples. Interestingly, in the Czech Republic 65.1% (1315/2019) of samples were identified as *C. andersoni* as well as in Ethiopia and Mongolia (Figure 4). A total of 13 other *Cryptosporidium* species were identified, including 64 cases of *C. hominis,* present in all continents with data except North and South America.

**Figure 4.**
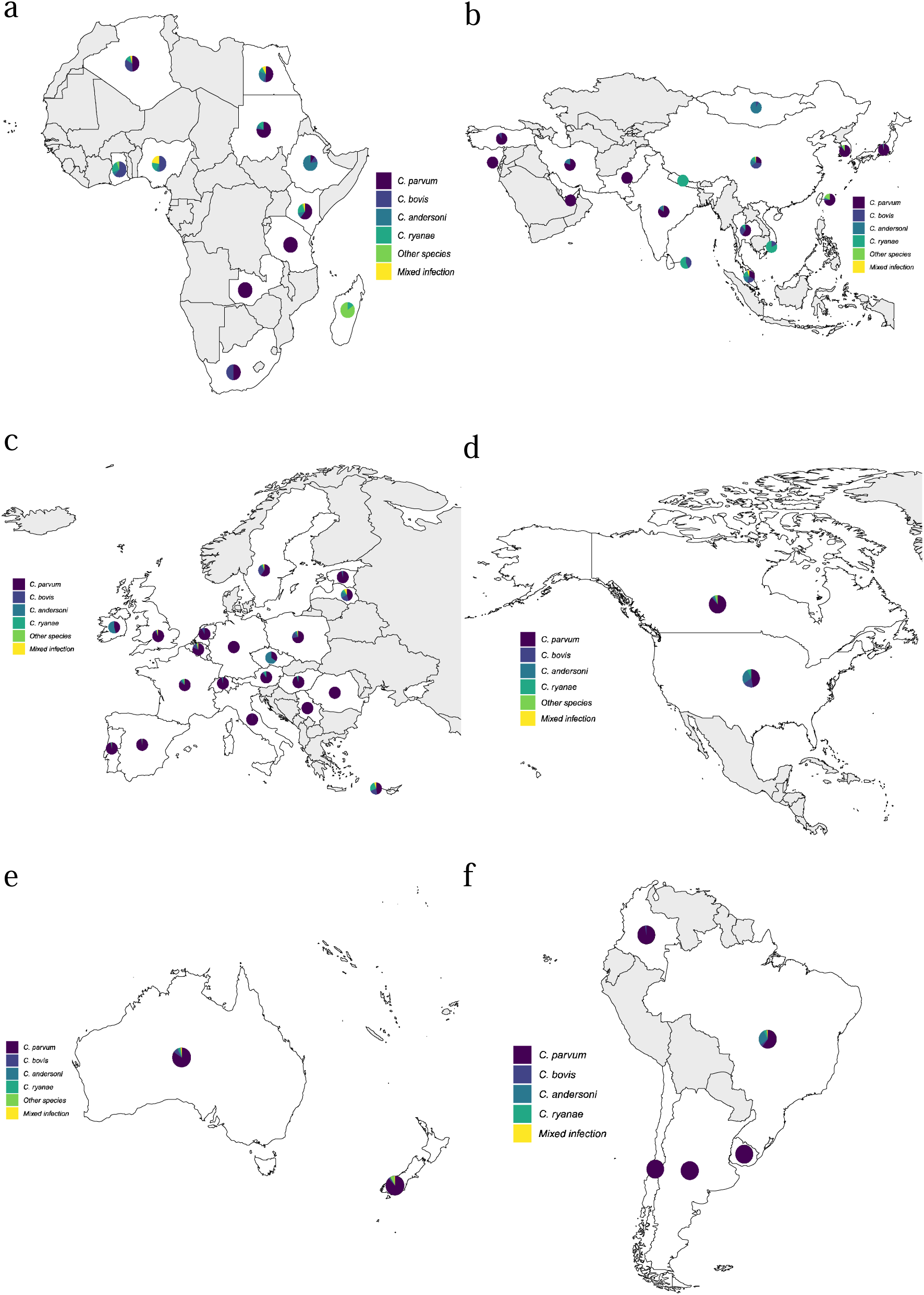
Distribution and frequency of *Cryptosporidium spp.* by continent.

For all studies reporting *C. parvum* cases, 34.0% (4311/12,664) were sub-typed, with three sub-type families identified: IIa (2991/4311; 69.4%), IId (1308/4311; 30.3%), and IIl (12/4311; 0.3%). The IIa family was found in all 6 continents with data, while the IId family was identified in all except North America and Oceania, and the IIl family was only found in Europe (Figure 5). The IIa family was the most prevalent one in all continents with data except Africa and Asia, where the IId family was most frequently identified, with the majority (1107/1308; 84.6%) of IId cases originating from China. For the 72 genotypes identified in the IIa family, 20 were identified once, 24 were identified 2-10 times, 20 were identified 11-99 times, and 8 were identified ≥100 times. The eight most common genotypes were IIaA15G2R1 (963/2991; 32.2%), IIaA18G3R1 (321/2991; 10.7%), IIaA16G1R1 (185/2991; 6.2%), IIaA13G2R1 (183/2991; 6.1%), IIaA17G2R1 (129/2911; 4.3%), IIaA19G2R1 (114/2991; 3.8%), IIaA19G3R1 (109/2991; 3.6%), and IIaA17G1R1 (100/2991; 3.3%). For the 24 identified IId genotypes, five were identified once, 15 were identified 2-20 times, and four were identified >100 times. The four most common genotypes were IIdA19G1 (514/1308; 39.3%), IIdA15G1 (292/1308; 22.3%), IIdA20G1 (290/1308; 22.2%), and IIdA14G1 (115/1308; 8.8%).

**Figure 5.**
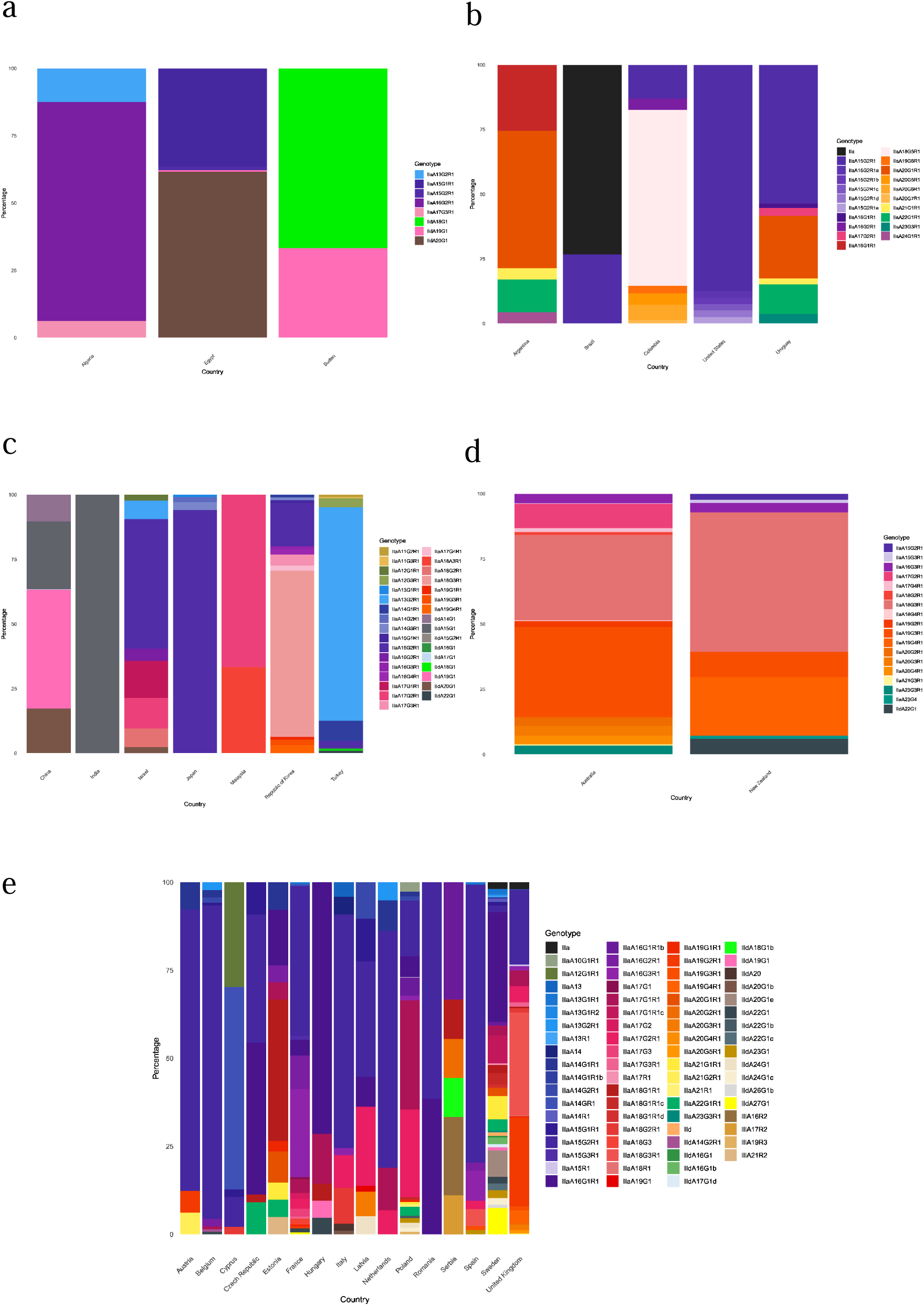
*C. parvum* genotypes distribution in different regions.

Publication bias was assessed using a funnel plot. The funnel plot shows obvious asymmetry with many datapoints falling outside of the funnel, which are both indicators of publication bias. This is consistent with the results of Egger’s test, indicating strong publication bias with p < 0.0001. There is also evidence of publication bias with p < 0.01 for all continents except North and South America where p < 0.05. By methodology, the results of the Egger’s test suggest publication bias (p < 0.0001) for CM, IFA, and PCR. Where studies indicated the keeping status of cattle (dairy or beef farms), Egger’s test also indicates publication bias with p < 0.01 (Table 1).

The meta-analysis revealed substantial heterogeneity across the included studies (*I* ^2^ = 99.4%, p < 0.0001). Potential sources of heterogeneity were explored using single and multiple predictor meta-regression across four categories: year of publication, detection method, study location (continent), and the diarrhoeal status of cattle (whether only diarrhoeic cattle were sampled in the study). Regarding single predictors, the publication year was a significant predictor (p = 0.017) but accounted for only 1.6% of heterogeneity. For diarrhoeal status, 87 of the included studies had clearly reported this data, and diarrhoeal status was a significant predictor amongst these studies (p = 0.0005), accounting for 11.6% of heterogeneity.

We then used a model selecting approach to identify the most suitable multiple-predictor model. When testing all four predictors, encompassing 87 studies with complete data, method was the predictor with the least significance (p = 0.2917). Refitting the model without the method predictor showed that the publication year was the least significant predictor (p = 0.2120). When only continent and diarrhoeal status were fitted, both predictors were significant (p = 0.0492 and p = 0.003, respectively), and accounted for 16.6% of heterogeneity. This model was the most suitable fit, however due to the limitations in available data it is not inclusive of all studies used in the meta-analysis.

## Discussion

This meta-analysis and systematic review identified a global pooled prevalence of 25.5% from 292 studies encompassing 144,523 cattle samples across 65 countries. Interestingly, when splitting publications by detection method, PCR and CM yielded an identical pooled prevalence of 25.0%, despite PCR being a more sensitive method [27]. CM is less sensitive, but cheaper and thus may still be capable of providing accurate estimates of *Cryptosporidium* prevalence in lower-income countries, which are the most underrepresented in this meta-analysis.

Cattle serve as a known reservoir of zoonotic cryptosporidiosis [20,28], with the majority of human cryptosporidiosis cases caused by either *C. hominis* or *C. parvum*. Additionally, the implications of cryptosporidiosis in cattle contributes to large financial losses, with the USDA in 2015 reporting losses of $3.87 billion due to cattle deaths from non-predator causes, of which 15.4% were due to digestive problems including cryptosporidiosis [29]. This, coupled with increasing global demand for livestock products to support the increasing population [30,31] makes monitoring the global distribution of *Cryptosporidium* in cattle vital. Concentrated Animal Feeding Operations (CAFOs) have been associated with a rise in zoonotic pathogens including *Cryptosporidium*, and the findings of this meta-analysis shows that a lower prevalence of *Cryptosporidium* is found in Africa and Asia, where cattle rearing is typically less intensive (Figure 1). Higher rates were found in Japan, Europe, and North America, which are countries with higher intensity rearing [17], with outbreaks reportedly rising in recent years in China due to increasing cattle CAFOs [32].

*C. parvum* is a known causative of diarrhoea in neonatal calves [10,11], and the majority of included studies sample this age group. A total of 20 of 36 countries with data report over 50% of samples being diarrhoeic, and *C. parvum* was the most frequent species identified in these countries (Figure 3, Figure 4). In the UK, it has been reported that costs associated with severe cryptosporidiosis symptoms in calves can amount to approximately £200 per calf when totalling veterinary costs and loss of market value due to weight loss [33], thus outbreaks of *Cryptosporidium* can lead to a significant financial burden for farms. Globally, there were also cases of cattle having diarrhoea that was not linked to a positive *Cryptosporidium* sample, indicating the potential presence other infectious agents frequently found in cattle as well as *Cryptosporidium* (Figure 3) [34].

The distribution of *Cryptosporidium* species was the most underreported in the African and South American continents. *C. parvum*, *C. bovis*, *C. andersoni*, and *C. ryanae* are the most common species infecting cattle [7,35], and this is shown in the meta-analysis, with *C. parvum* being the most frequently identified species, likely due to the majority of included studies involving neonatal calves (*C. bovis* was the most frequently identified species in three countries, *C. andersoni* in four countries, *C. ryanae* in one country, and *C. xiaoi* in one country). All five of the most frequent species reported in humans (*C. hominis*, *C. parvum*, *C. meleagridis*, *C. canis*, and *C. felis*) were identified [28], furthering the evidence of cattle acting as a zoonotic reservoir. Not only is the global *C. parvum* distribution capable of compromising the health of calves, but the identification of species capable of infecting humans also poses a public health risk. Global estimates suggest that the yearly oocyst load of livestock manure is 3.23 x 10^23^, with cattle being large contributors [36]. Contamination of food and water sources with oocysts shed by cattle has been associated with foodborne [37–39] and waterborne [39–41] outbreaks.

The *C. parvum* IIa subfamily was the most widely distributed one, making up the majority of infections in Europe, North America, Oceania, and South America, while the IId subfamily was the source of most infections in Africa and Asia (Figure 5). North America was significantly under-reported, with 2.2% of *C. parvum* cases genotyped, while the five other continents had genotyped between 36.4-46.3% of cases. There was also large variation in the number of different genotypes identified in each country, particularly noticeable in Sweden, where 32 different genotypes were found. The highly transmissible IIaA15G2R1 [5,14] was the dominant IIa subfamily genotype, particularly in Europe and South America. This genotype is also the dominant in human *C. parvum* infections in countries where it is a common bovine genotype [5,39]. The IIaA18G3R1 genotype was also frequently identified in Europe and Oceania and has been the cause of human infections in Ireland [42]. The dominant IId subfamily genotype, IIdA19G1, was identified primarily in cattle from China, with only several cases in Africa and Europe. This genotype has been linked to human infections in China [43,44], and Scotland [45].

Our meta-analysis aims to provide a recent estimate of global *Cryptosporidium* prevalence in cattle using studies published in the last 20 years. This search strategy gave a large sample size of 144,523 cattle across 6 continents and 65 countries. However, despite this large dataset, there are several limitations to our study. First, only data of publicly accessible journals and published in English are included. Additionally, there are still many countries with no published data, and for some included countries there were few publications available meaning many regions were underrepresented. This became evident particularly when quantifying *Cryptosporidium spp.* and *C. parvum* genotype distribution, where the number of species or genotypes identified from positive samples varied greatly between continents and countries. Statistical analysis also revealed high levels of publication bias and heterogeneity, which can affect the results (Table 1) [46]. This was explored using meta-regression, and the publication year and diarrhoeal status of the sampled cattle were identified as potential sources of heterogeneity as single predictors, however there are still further factors that could be explored such as the farming status, rearing intensity, or sex of sampled cattle, but this data is not frequently provided in detail in publications. We show that the diarrhoeal status and continent resulted in the best fitting multiple meta-regression model, accounting for 16.6% of heterogeneity in 87 studies. Given that the majority of studies did not have complete data for these predictors, the amount of heterogeneity accounted for by the tested predictors remains unknown across the entire dataset. However, the study continent and the diarrhoeal status of sampled cattle are likely sources of heterogeneity due to sampling bias created by only sampling symptomatic cattle, and through different continents having differing prevalence and reporting rates of *Cryptosporidium*. Given the limitations from the lack of predictor data, it is evident that further work is needed to ensure equal and accurate reporting of global *Cryptosporidium* prevalence, especially regarding the public health risk posed by cryptosporidiosis and zoonotic species of *Cryptosporidium*.

## Conclusions

The findings of this meta-analysis suggest widespread bovine *Cryptosporidium* infection, with approximately one in four sampled cattle testing positive. The data also shows significant variance by continent and country, although the underrepresentation of certain regions is a likely contributor to this. Regions with higher numbers of CAFOs also had higher infection rates. *C. parvum* was the most identified *Cryptosporidium* species, with the IIa subfamily found globally and the IId subfamily most frequently found in China. The prevalence of diarrhoea in sampled cattle also varied, however in countries with higher rates of cattle exhibiting diarrhoea, *C. parvum* was the dominant species. Effective control measures should continue to be developed and implemented to prevent infection in cattle, which can limit the risk of transfer to humans. Additionally, further efforts should be undertaken to report on the prevalence and transmission in countries with limited or no data to ensure that control measures are implemented effectively where they are needed. To that end, adopting a ‘One Health’ approach, which underscores the interconnectedness of human, animal, and environmental health, is crucial for developing comprehensive strategies to control and prevent *Cryptosporidium* infections, thereby safeguarding public health and reducing economic losses in the livestock industry.

## Abbreviations

CM: Conventional Microscopy
ELISA: Enzyme-linked Immunosorbent Assay
ICT: Immunochromatographic Test
IFA: Immunofluorescence Antibody Test
PCR: Polymerase Chain Reaction

## Notes

### Competing Interest Statement

The authors have declared no competing interest.

